# Cluster Tendency Assessment in Neuronal Spike Data

**DOI:** 10.1101/285064

**Authors:** Sara Mahallati, James C. Bezdek, Milos R. Popovic, Taufik A. Valiante

**Affiliations:** Institute of Biomaterials and Biomedical Engineering, University of Toronto, Canada; Toronto Rehabilitation Institute, University Health Network, Toronto; Krembil Research Institute, University Health Network, Toronto; Computer Science and Information Systems Departments, University of Melbourne, Australia; Division of Neurosurgery, University of Toronto

**Keywords:** spike sorting, single units, cluster assessment, unsupervised learning, dimensionality reduction, t-SNE, iVAT, Dunn’s Index

## Abstract

Sorting spikes from extracellular recording into clusters associated with distinct single units (putative neurons) is a fundamental step in analyzing neuronal populations. Such spike sorting is intrinsically unsupervised, as the number of neurons are not known a priori. Therefor, any spike sorting is an unsupervised learning problem that requires either of the two approaches: specification of a fixed value c for the number of clusters to seek, or generation of candidate partitions for several possible values of c, followed by selection of a best candidate based on various post-clustering validation criteria. In this paper, we investigate the first approach and evaluate the utility of several methods for providing lower dimensional visualization of the cluster structure and on subsequent spike clustering. We also introduce a visualization technique called improved visual assessment of cluster tendency (iVAT) to estimate possible cluster structures in data without the need for dimensionality reduction. Experimental results are conducted on two datasets with ground truth labels. In data with a relatively small number of clusters, iVAT is beneficial in estimating the number of clusters to inform the initialization of clustering algorithms. With larger numbers of clusters, iVAT gives a useful estimate of the coarse cluster structure but sometimes fails to indicate the presumptive number of clusters. We show that noise associated with recording extracellular neuronal potentials can disrupt computational clustering schemes, highlighting the benefit of probabilistic clustering models. Our results show that t-Distributed Stochastic Neighbor Embedding (t-SNE) provides representations of the data that yield more accurate visualization of potential cluster structure to inform the clustering stage. Moreover, The clusters obtained using t-SNE features were more reliable than the clusters obtained using the other methods, which indicates that t-SNE can potentially be used for both visualization and to extract features to be used by any clustering algorithm.

## 1 Introduction

Recording of extracellular signatures of action potentials, referred to as spikes, is a standard tool for revealing the activity of populations of individual neurons (single units). Single unit activity contains fundamental information for understanding brain microcircuit function in vivo and in vitro [Buzsáki, 2004]. Inferences about network activity can be made by identifying coincident activity and other temporal relationships among spiking patterns of different neurons [Brown et al., 2004]. However, the reliability of these inferences is strongly influenced by the quality of spike sorting, i.e., the detection and classification of spike events from the raw extracellular traces with the goal of identifying single-unit spike trains. Poor sorting quality results in biased cross-correlation estimates of the spiking activity of the different identified units [Ventura and Gerkin, 2012].

The typical workflow for spike sorting includes spike detection, feature extraction, and clustering. While detection is pretty straightforward and can be efficiently done with simple thresholding, the feature extraction and clustering procedures are far from being satisfactorily settled [Rossant et al., 2016]. It has been estimated that single or tetrode type electrodes (i.e. impedance < 100*K*Ω) can record neuronal activity within a spherical volume of 50 *μ*m radius with amplitudes large enough to be detected (> 60*μ*V). This volume of brain tissue constitutes about 100 neurons. While noting that many neurons are expected to be silent [Buzsáki, 2004, Shoham et al., 2006], commonly, not more than a handful identified neurons are reported per electrode. Studies on current sorting algorithms used for individual electrode recordings have shown that they are limited in distinguishing 8 to 10 out of 20 units with less than 50% false positive and false negative rates [Niediek et al., 2016, Pedreira et al., 2012]. Other methods using high density electrode arrays reported simulations with no more than 10 units [Pachitariu et al., 2016, Yger et al., 2016].

Since we can’t physiologically verify how many neurons have been recorded, assigning the spikes within a recording to individual neurons remains a fundamental technical issue. The sorting is in essence an unsupervised learning challenge. Therefore, methods require one of two approaches: specification of a fixed value of the number of clusters to seek (*c*); or generation of candidate partitions for several possible values of c, followed by selection of a best candidate based on various post-clustering validation criteria. Moreover, improving spike classification to correctly identify cell types is a topic of interest highlighted by initiatives that aim to characterize and reconstruct different cell types in the brain and their role in health and disease [Jorgenson et al., 2015, Markram et al., 2015]. For that goal, Armañanzas and Ascoli [2015] list the identification of the number of clusters as the first outstanding question in techniques for neuronal classification.

In summary, identifying the spike trains of individual units within a recording is a three-faceted problem: (i) assessing the cluster tendency in the pre-clustering phase (before initializing any clustering algorithm); (ii) clustering (i.e. finding partitions of the data); and (iii) evaluation of the validity of the clusters that have been isolated, post-clustering [Bezdek, 2017]. Spike sorting algorithms usually start by projecting the data to a lower dimensional space. There are several reasons to do this. For example, lower dimensional data usually reduces such as reduction of computation time. In this paper, the fundamental reason for dimensionality reduction (essentially to two or three dimensions) is that 2D and 3D projections allow visualization of high dimensional input data. In turn, this facilitates the choice of a few selected values of the integer c. Pre-specification of c is needed by almost all clustering algorithms as an input parameter (hyper parameter). In algorithms such as density based clustering or mean shift, the choice of c is implicit in the choice of parameters such as *γ* (cluster density times cluster distance) or *h* (the bandwidth parameter), respectively. In practice, since reduced dimensionality embedding of the data often does not provide visually well separated clusters, it is common to exclude large number of spikes and only take into account a small core portion of the subsets that seems to have well-isolated clusters [Dehghani et al., 2016]. Omitting spikes to obtain well-separated clusters may lead to single units with recognizable spike waveforms, but it discards spikes that, as mentioned before, are fundamental for analyses of temporal structure of spiking activity [Cohen and Kohn, 2011, Pazienti and Grün, 2006]. Therefore, the bottleneck in spike sorting is at the pre-clustering stage: viz., inaccuracy of the assumed data structure that is inferred by visualization of it in the lower dimensional space. If clustering is to be done in a lower dimensional data space, errors here will affect both the initial estimate of the cluster number and the performance of the clustering algorithm. Thus, this study concerns itself with visual assessment in the pre-clustering stage.

We compare the visualization of neuronal spike data created using six methods (i) three well-known dimensionality reduction techniques: principle component analysis (PCA), t-student stochastic neighborhood embedding (t-SNE) and Sammon’s algorithm, (ii) two methods that extract features from the waveforms: wavelet decomposition and features such as peak to valley amplitude and Energy (PV-E), and (iii) a method that operates directly on waveforms in the input space: improved visual assessment of tendency (iVAT). The analysis is performed on two different types of ground truth data (labeled data): simulated spike sets and real recorded spike sets, called dataset-1 and dataset-2, respectively. Our results indicate that iVAT often suggests a most reasonable estimate for the primary (or coarse) cluster structure, while t-SNE is often capable of displaying finer cluster structure. While iVAT is only a visualization tool, t-SNE can be used for both visualization and to extract features. We provide an objective measure of comparison between t-SNE and the other methods, we evaluate the quality of partitions obtained by clustering in the upspace (input dimensional space; i.e., the waveforms) and also in the five two-dimensional representations. This test is performed by running k-means and generating a number of clusters equal to the actua (i.e. known) number of units. The quality of the partitions generated by each method is evaluated with *Dunn’s index (DI)* (an internal index describing the intrinsic quality of the generated clusters); the generalized Dunn’s index GDI_33_; and the *Adjusted Rand’s index (ARI)* (an external measure of agreement between computed partitions and the ground truth partition).

The outline of the paper follows: Section 2 describes the datasets that we used (2.1), defines the problem (2.2), explains the methods used for data visualization (2.3 and 2.4), and lastly describes the measures used for evaluating clustering structure of the data (2.5). The results of the experiments on the datasets are given in section 3. Insights gained from the experiments are summarized in section 4.

## 2 Materials and methods

### 2.1 Datasets

The importance of model data or ground truth data, where the label or membership of each spike to an individual neuron is known, for spike sorting validation is emphasized by Einevoll et al. [2012]. We use two labeled datasets: our first experiment uses simulations of extracellular traces as model data or surrogate ground truth data (hereafter called dataset-1), and the second experiment uses data obtained from in-vivo experiments as real ground truth data (hereafter called dataset-2).

#### Dataset-1

Pedreira et al. [2012] simulated exracellular traces that contain the activity of 2 to 20 neurons with additive background noise. The single unit spike activity is generated by using average spike waveforms (templates) compiled from Basal Ganglia and Neocortical recordings. The background noise (i.e., LFP noise) is simulated by superimposition of thousands of spikes at random times which were then scaled down to a standard deviation of 0.1. Each simulated trace also contains multi-unit activity, which was generated by superimposing 20 waveforms with normalized amplitudes limited to 0.5. Dataset-1 thus provides us with simulated extracellular traces each containing 3 to 21 subsets of spikes. For each cluster number *c*, five simulations using different sets of templates were generated (for a total of 95 datasets). The spikes were detected by voltage thresholding. The length of each spike waveform is 2 ms, with a sampling rate of 24 kHz, comprising 48 sample points. Thus, the upspace dimension for subsets in Dataset-1 is 48.

#### Dataset-2

We used the in vivo simultaneous intracellular and extracellular hippocampal recording datasets that are publicly available from the CRCNS website [Henze et al., 2009]. These are raw data of simultaneous intracellular and extracellular recordings from neurons in the CA1 region of the hippocampus of anesthetized rats. The experimental procedure consisted of inserting extracellular electrodes (either tetrodes, 13-*μm* polyimide-coated nichrome wires, or a single 60-*μm* wire) into the cell body layer of CA1, confirming the presence of unit activity in the recordings, and then inserting intracellular sharp-electrodes into the same region in close proximity to the extracellular electrode to impale a single neuron and record stable action potentials induced by current injections. With this method, it was possible to capture simultaneous spikes in the intracellular and extracellular recordings. We detected the intracellular spikes, and used those spike times to extract the extracellular spike train of that neuron (i.e. a labeled subset of spikes). Each spike waveform is 2 ms long which, with a sampling rate of 40 kHz, results in 80 sample points, so the upspace dimension of each spike and subsequent sets in dataset-2 is 80. Dataset-2 is valuable because each spike waveform in the subset is a recorded signal from a physiological setting; hence, the variability in the probability distribution of each subset comes from either natural (e.g. the effect of other current sources in the extracellular medium) or experimental conditions (e.g. electrode drift). Each recorded trace has one labeled subset of spikes. Hence, to generate each mixture, we used the extracted spike subsets from different traces. In total, we obtained 9 subsets of spikes from the database and then used combinations of 2,3,4, … to 9 of these subsets to create datasets containing spikes of 2 or more neurons (for a total of 502 datasets).

In summary, the data for our experiments were mixtures of subets of spikes each with different population size in either 48 or 80 dimensional input space.

### 2.2 Problem definition

Let 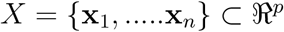 denote a set of vector data representing *n* spikes generated by one or multiple neurons. The coordinates of **x**_*i*_ are voltage samples that describe a spike event (they are always voltage samples in this article). The non-degenerate crisp *c*-partitions of the *n* objects in a set *X* can be represented by a *c × n* matrix *U* in *M*_*hcn*_, written in terms of the *c* crisp subsets of it (the clusters X_*i*_) as

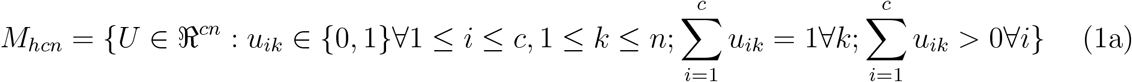

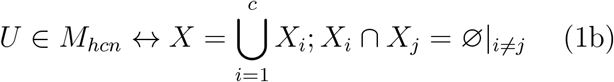

Finding clusters in *X* comprises three steps: deciding how many clusters (*c*) to look for; constructing a set of candidate partitions {*U* ∈ *M*_*hcn*_} of X; and selecting a “best” partition from CP (cf. equation (2) below) using a *cluster validity index (CVI)*.

### 2.3 Dimensionality reduction and feature extraction

Data vectors in 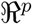 usually have high dimensionality (*p* > 3) (e.g., images, videos, and multi-variate data streams). Feature selection and dimensionality reduction algorithms are used to (i) make pre-clustering visual assessment of the structure of the data and (ii) to improve the performance of data-driven procedures, such as those for classification and clustering. Typical approaches for these procedures include those discussed in Zhao et al. [2013] and van der Maaten et al. [2009]. Methods for obtaining an “optimal” set of features (the intrinsic dimension of the input data) for clustering abound. Campadelli et al. [2015] have a good survey of such methods. We point out finding the intrinsic dimension of a data set is not directed towards visual assessment, since the intrinsic dimension (there are many competing algorithms that find different features) is usually greater than 2 or 3.

Here we focus on visualization based on several well-known dimensionality reduction algorithms that have been used in a multitude of domains including neurosciences; we refer the interested reader to van der Maaten et al. [2009] and references therein for technical details.

Principal component analysis (PCA) is one of the most important and widely utilized linear dimensionality reduction techniques [Theodoridis, 2009]. In order to find a low-dimensional subspace that accounts for the maximum amount of projected variance, PCA projects the data along the directions given by the leading eigenvectors of the data covariance matrix, i.e., the directions associated with the largest eigenvalues of the sample covariance matrix.

In neuroscience research, another common approach is to extract features of the waveforms that have a physical meaning such as waveform’s negative or positive amplitudes, known as valley and peak, their ratio or width, or waveform’s energy, among others [Hattori et al., 2015, Truccolo et al., 2011]. For our experiments we selected peak to valley (PV), and energy (E), hereby called PV-E. We remark that the PV-E features contain essentially the same information as the min peak-max peak features used by [Takahashi et al., 2003]. Another method based on wavelet transforms that enables visualizing the data in the wavelet coefficient subspace has also been successfully implemented in clustering packages such as Waveclus and Combinato [Niediek et al., 2016, Quiroga et al., 2004].

We also consider two nonlinear dimensionality reduction techniques. The first of these is t-SNE (t-student Stochastic Neighbor Embedding), developed by van der Maaten and Hinton [2008]. It works by converting Euclidean distances between high-dimensional input data into conditional probabilities. In doing so, t-SNE converts the geometric notion of similarity into a statistical concept: if *x*_*j*_ is a neighbor of *x*_*i*_, then the conditional probability *p*_*j|i*_ is high. Then, t-SNE finds low-dimensional representations *y*_*i*_ and *y*_*j*_ of *x*_*i*_ and *x*_*j*_ by minimizing the discrepancy between the upspace *p*_*j|i*_ and downspace conditional probabilities *q*_*j|i*_, technically achieved by minimizing the Kullback-Leibler divergence between them. The objective of t-SNE is to minimize the sum of the divergences over all the data points. The downspace dimension is a choice made by the user.

Two features of t-SNE should be noted. First, it is not a linear projection like PCA but rather has a non-convex cost function, so its output may be different for different initializations. Second, it is a parametric technique. Different settings of hyperparameters such as the learning rate, the *perplexity*, and the iteration rate in the t-SNE algorithm generate different maps in the scatterplots, and may cause misinterpretation of the data structure [van der Maaten and Hinton, 2008].

The main parameter that affects the results of t-SNE is the perplexity, which is the limiting condition for the entropy of the probability distribution of the similarities of datapoints in the upspace. This means that the variance of the Gaussian that is centered over each datapoint, i.e., the extent of the neighborhood around that point, is limited by the choice of perplexity.

This limitation affects each datapoint separately based on the local density of the data. This is the feature that enables t-SNE to avoid crowding points in the center of the map so that cluster structure of the data in the upspace data is often seen in the t-SNE downspace projection. This feature, however, comes at the cost of sacrificing the shape of the distribution so that the distances between the clusters may not be meaningful. In other words, it is not possible to infer reliable spatial information from the topology of the low-dimensional maps.

Fortunately, the topology is not relevant for our application: viz. suggesting clusters in the neuronal waveform data. The optimal choice of perplexity is dependent on the number of points (spikes) in the dataset. We found that for neuronal datasets with thousands of spikes (data points), as long as the extreme values in the parameter ranges are not selected, the t-SNE algorithm is not very sensitive to changes in perplexity. On the other hand, the reliability of t-SNE visualizations seems to decrease as the number of samples decreases. See [Mahallati et al., 2018a] for an example.

We also consider another traditional nonlinear dimensionality reduction technique called the Sammon mapping [Sammon, 1969], which is one form of multidimensional scaling. Multidimensional scaling (MDS) seeks a low dimensional embedding of the input data while preserving all pairwise Euclidean distances (In a more general setting, t-SNE can be interpreted as a form of probabilistic MDS). However, high-dimensional data usually lies on a low-dimensional curved manifold, such as in the case of the Swiss roll [Tenenbaum et al., 2000]. In such cases, preserving pairwise Euclidean distances will not capture the actual neighboring relationships: the actual distance between two points over the manifold might be much larger than the distance measured by the length of a straight line connecting them, i.e., their Euclidean distance). Sammon mapping improves upon classic multidimensional scaling by directly modifying its original cost function, i.e., the distortion measure to be minimized. In particular, the Sammon mapping cost function weights the contribution of each pair of data points relative to the overall cost by taking into account the inverse of their pairwise distance in the original high-dimensional input space. In this way, Sammon mapping often preserves the local structure of the data better than classical multidimensional scaling.

While these five methods do not all produce lower dimensional data with an analytic projection function, we will call all downspace data sets projections.

### 2.4 iVAT

There are a number of imaging techniques that can be applied directly to the upspace data before clustering it. Here we describe the iVAT method described in [Havens and Bezdek, 2012], which is a generalization of the original VAT algorithm given by [Bezdek and Hathaway, 2002]. *Improved Visual Assessment of Tendency* (iVAT) is a visualization tool that uses any dissimilarity matrix, D, of the data to display potential cluster structure. The steps of the iVAT method are the following. The vectors in the dataset are represented as vertices in a complete graph, with the distances between them the weights of the graph. The algorithm first finds the longest edge in the graph. Then, starting at either end, it finds the minimal spanning tree (MST) of D based on Prim’s algorithm. Then, it reorders the rows (and columns) of D based on the order of edge insertion in the MST, creating D* (up to this point this is the original VAT algorithm). Then, iVAT transforms D* to D’* by replacing each distance *d*_*ij*_ in D* with the maximum edge length in the set of paths in the MST between vertices *i* and *j*. When displayed as a gray-scale image, I(D’*), possible clusters are seen as dark blocks along the diagonal of the image. Images of this type are often called *cluster heat maps* in the neuroscience literature.

iVAT is not a clustering method or measure of performance for clustering algorithms, but a method to visually extract some information about the cluster structure from the input space before application of any clustering algorithm. iVAT does not alter the physical meaning of the input data (even after the shortest path transformation), it just rearranges the objects in a way that emphasizes possible cluster substructure. The recursive computation of *D‣\** given in Havens and Bezdek [2012] is *O*(*n*^2^). Appendix A.2 contains the pseudocode for iVAT. The iVAT algorithm requires no parameters to pick other than the dissimilarity function (d) used to convert X to D. This input matrix can actually be a bit more general than a true distance because its only requirements are that *D* = *D*^*T*^; *d*_*ij*_ ≥ 0∀*i*, *j*; *d*_*ii*_ = 0∀*i*. The most important points about this display technique are that it is applied directly to (a distance matrix of) the upspace data, so there is no distortion of the structural information introduced by a feature extraction function from the upspace to a chosen downspace, and iVAT preserves the physical meaning of the measured features. While any vector norm can be used to build an input matrix D(X) from a set X of feature vectors, the only distance used in this article is Euclidean distance. It is very important to understand that an iVAT image merely suggests that the input data has a certain number of clusters. Since iVAT can produce images from data of arbitrary dimensions, we can use it (or its scalable relative siVAT, Kumar et al. [2017]) to make a visual estimate of possible cluster structure in any upspace. While the iVAT algorithm is occasionally “wrong” (misleading), iVAT images usually provide some idea about the cluster structure of the input data [Bezdek, 2017].

Thus, iVAT provides clues about potential starting points for finding a useful partition of the input data. Mahallati et al. [2018a] have shown the connection of VAT and iVAT to Dunn’s index and single linkage (SL) clustering. The intensity of the blocks in iVAT images are a (more or less) visual representation of the structure identified by single linkage clustering for labeled or unlabeled data. This suggests that iVAT might be regarded as a tool for “taking a peek” at a specific type of data structure in the input space.

### 2.5 Evaluating cluster quality

Cluster validity comprises computational models and algorithms that identify a “best” member amongst a set of *candidate partitions (CP)*

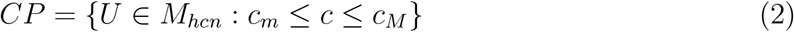

where *c*_*m*_ and *c*_*M*_ are the minimum and maximum specified values of the numbers of clusters sought.

The approach to identify a “best” partition U (and concomitant value of *c*) in *CP* can be internal: using only information from the output results of the clustering algorithm, or external: using the internal information together with an outside reference matrix, usually the ground truth labels. Here, we use a classic internal scalar measure called *Dunn’s index* (DI) [Dunn, 1973], and one generalization of it given by Bezdek and Pal [1998] called the generalized Dunn’s index (GDI_33_). Dunn defined the diameter of a subset *X*_*k*_ as the maximum distance between any two points in that subset (Δ(*X*_*k*_)), and the distance between subsets *X*_*i*_ and *X*_*j*_ as the minimum distance between any two points of the two subsets (*δ*(*X*_*ij*_)). This index is based on the geometrical premise that “good” sets of clusters are compact (dense about their means) and well separated from each other. Larger values of DI imply better clusters, so we call DI a max-optimal cluster validity index (CVI).

Let *X*_*i*_ and *X*_*j*_ be non empty subsets of 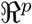, and let 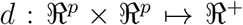 be any metric on 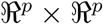. Define the diameter Δ of *X*_*k*_ and the set distance *δ* between *X*_*i*_ and *X*_*j*_ as:

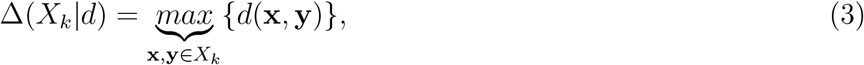

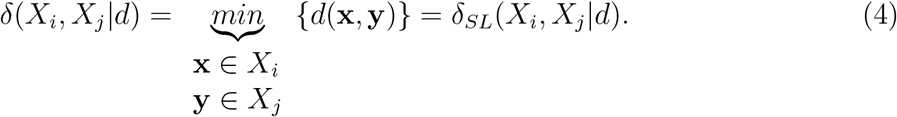

Then for any partition *U ↔ X* = *X*_1_ ∪ … *X*_*i*_ ∪ …*X*_*c*_, **Dunn’s separation index** of *U* is:

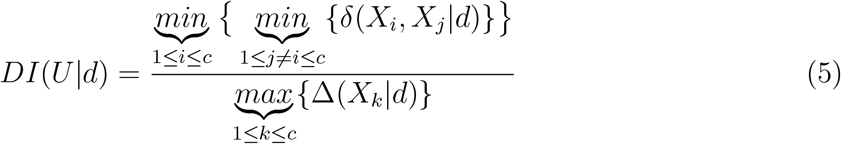

Since we have labeled mixtures, we can calculate Dunn’s index on ground truth partitions in the upspace (input dimensional space) to give a measure of the compactness and isolation quality of the subsets in the original space. We have previously shown that this measure usually correlates with the quality of the visual assessment of potential cluster structure given by iVAT [Mahallati et al., 2018a]. In the present work, we will also use a generalized version of Dunn’s index developed by Bezdek and Pal [1998] that alters the average distance from the mean as Δ and the average linkage clustering distance as *δ*. The **Generalized Dunn’s index** (*GDI*_33_)is:

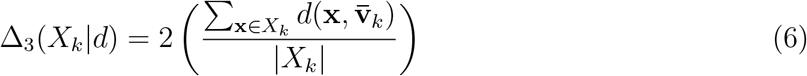

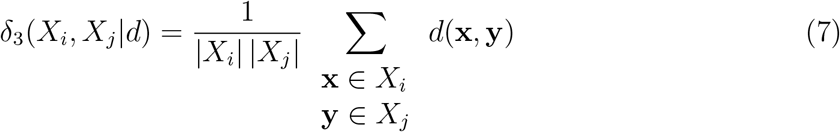

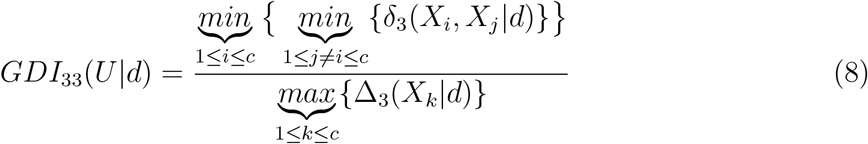

where 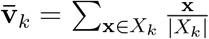 is the mean or centroid of the cluster. The notation |*d* in equation 3 to 8 for Δ and *δ* indicate that these formulas are valid for any metric d on the input space.

It has been shown that *GDI*_33_ is more robust with regards to sensitivity to outliers and hence produces more meaningful values for real life datasets with abundant aberrant points [Arbelaitz et al., 2013].

To evaluate the quality of the different clustering approaches we used the external **adjusted Rand index (ARI)** developed by Hubert and Arabie [1985], which is a well-known and fairly reliable criterion for performance assessment of the clustering results. Let *V* ∈ *M*_*hrn*_ be the crisp partition of the n objects possessing r clusters, according to ground truth labels. Let *U* ∈ *M*_*hcn*_ be any crisp partition of n objects with the c clusters generated by any clustering algorithm. Note that r does not necessarily equal c. The ARI is a measure of similarity between U and V, computed as:

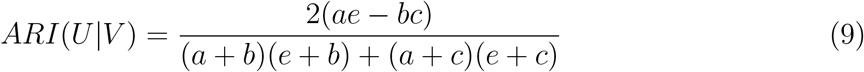

where,

- a = Number of pairs of data objects belonging to the same subset in U and V.
- b = Number of pairs belonging to the same subset in V but to different subsets in U
- c = Number of pairs belonging to the same subset in U but to different subsets in V.
- e = Number of pairs not in the same subset in V nor the same subset in U.

Hubert and Arabie [1985] developed this correction to eliminate bias due to chance from Rand’s index. The ARI is also a max-optimal index in the sense that larger values imply a better match between the ground truth and the results of the clustering. As evident from the equation, this formula incorporates simpler measures such as percentage of spikes in the real set present in the sorted set or percentage of spikes in the sorted set that come from the real set.

## 3 Results and discussion

### 3.1 Visual assessment of cluster tendency

It is impossible to make a direct visual assessment of a set of recorded spike waveforms 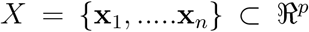, since each waveform has more than three voltage samples (i.e., dimensions), *p* > 3. The *upspace* dataset X can be mapped to a *downspace* dataset 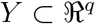 by a feature extraction function 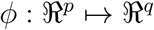 in many different ways. Dimensionality reduction methods are commonly employed for visualization purposes to gain insights into the data structure; and to provide clustering algorithms with lower-dimensional data to increase the computational efficiency. Next, we will demonstrate that different dimensionality reduction methods provide different scatterplots of the data, and hence, visually suggest different numbers of clusters. Towards this end, we used spike subsets of the simulated dataset that includes 5 simulations for each combination of different number of spike subsets for c from 3 to 21. Below we show some representative results: two cases of c=3 (one with low Dunn’s index, DI, and one with higher DI) and then one case each for c=5, c=10, c=15 and c=20.

Figure 1 shows two cases from the dataset with c=3. The colors in Figure 1 correspond to the three data labels. Bear in mind, as you view this and subsequent figures, that in the real case, the data are always unlabeled, so the projected data will be just one color, and the apparent structure will be much less evident than it seems to be in these figures. Figure 1(a) is a ‘good’ case in which all the algorithms map the spikes to projections with visually well-separated clusters and iVAT agrees with them (the larger diagonal block contains two less apparent,sub-blocks). In 1(b) however, all 2D projections except t-SNE produce a single cluster (when plotted without colors), while t-SNE seems most successful in separating the three subsets (arguably, t-SNE shows c=2 clusters when colors are omitted). The iVAT image suggests c=2, conforming to the apparent (uncolored) pair of t-SNE clusters. The low value of DI is a warning that there is not much separation between these three subsets of waveforms.

Figures 2 to 5 show representative mixtures of c=5, 10, 15 and 20 component mixtures. These examples, and many others not reported here, show that iVAT and t-SNE usually provide useful visual estimates of the number of clusters up to around c=15, but the other methods almost always fail with c = 4 or more subsets. We had 5 cases for each number of subsets (e.g. 5 different cases of mixture at c=10, etc.) and overall t-SNE provided the most consistent estimate of the presumptive numbers of mixture components. There were some cases for which iVAT failed to display the expected number of dark blocks in mixtures having fewer than 10 components. The block structure in some of the reproduced iVAT images is more apparent at higher resolutions than shown here. Our experiments suggest that iVAT is somewhat sensitive to noise in the waveforms, which often manifests itself as a falloff in intensity towards one end of the diagonal. See Figure 2 for an example.

**Figure 1:**
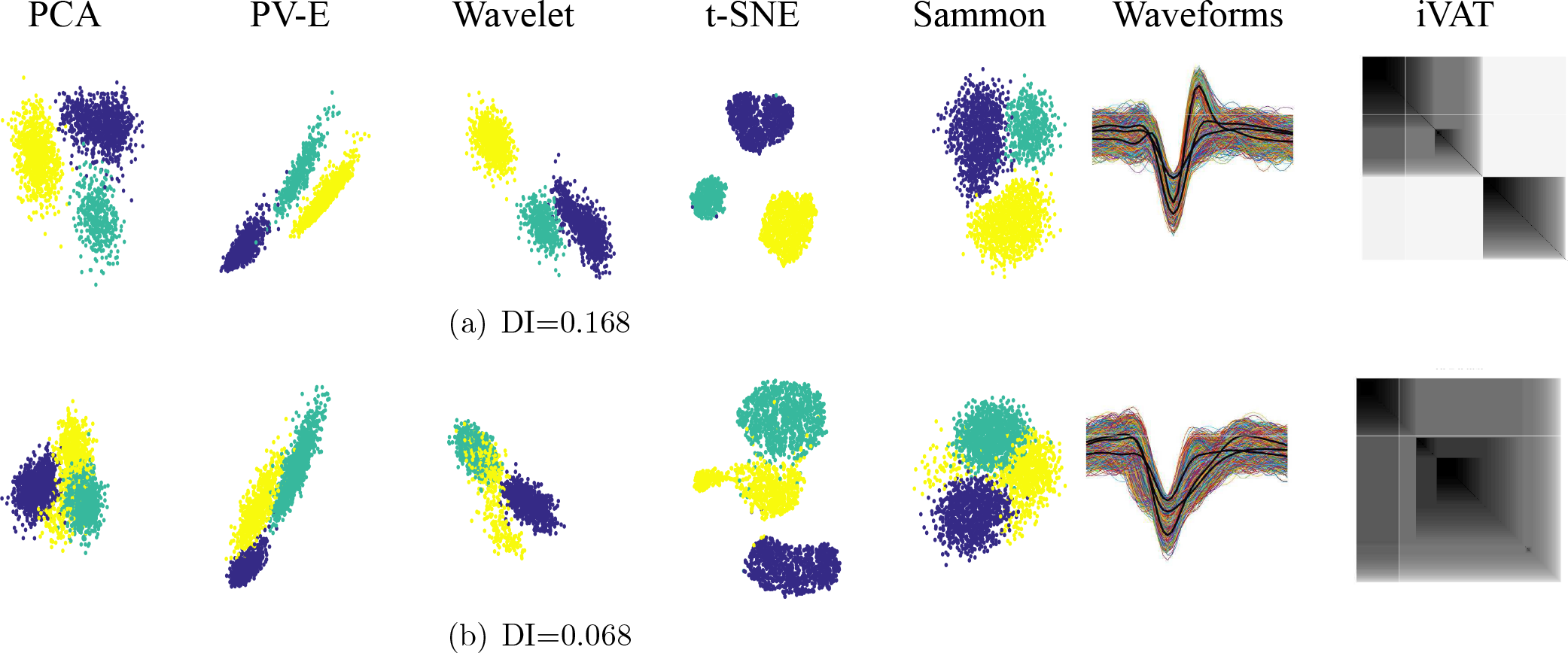
Two different simulations at c=3

Overall, from the t-SNE visualizations of the data, we infer that there is a common waveform shape among many cells suggested by the existence of a large central region populated by most of the clusters. However, there are cells that have clearly distinct signatures. The individual clusters exhibit variability: for some neurons the spike waveforms define homogeneous and compact clusters, while others are elongated clusters in the nonlinear space. This suggests that the relation between waveform samples for different neurons is different (or the way the waveform samples interrelate is different in different neurons’ spikes).

**Figure 2:**
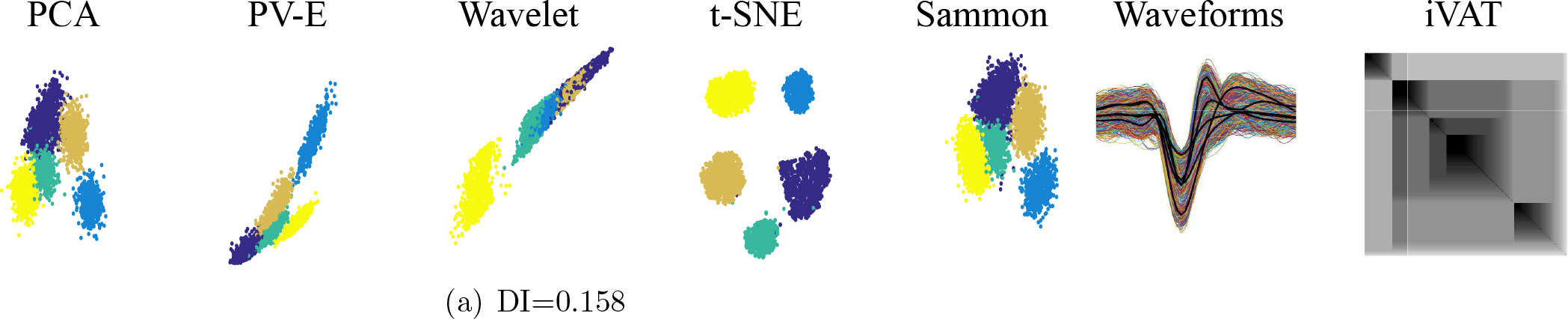
Simulation at c=5. The iVAT image displays 4 clear blocks and some disconnected data due to noise in the lower right: the only projection that clearly shows c=5 is t-SNE

**Figure 3:**
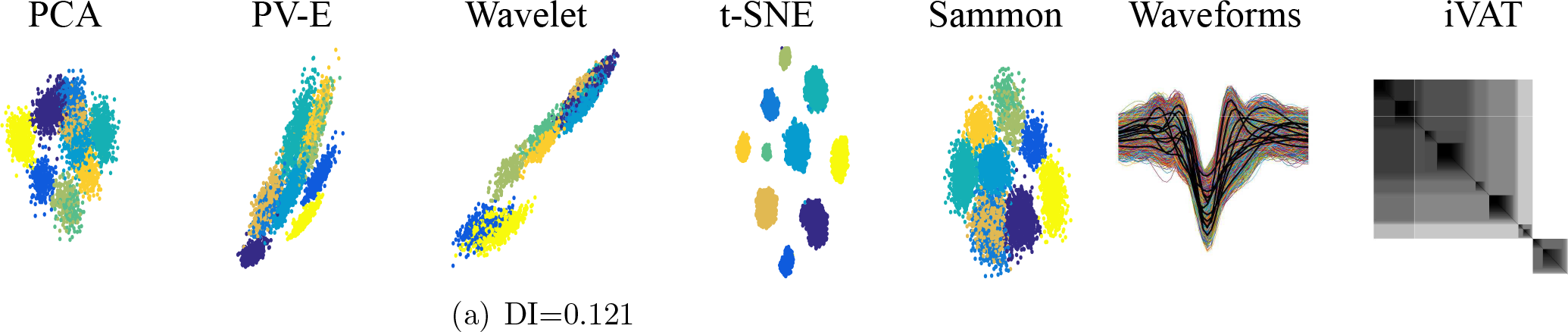
Simulation at c=10. The iVAT image displays 10 blocks; the projection that clearly shows c=10 is t-SNE

**Figure 4:**
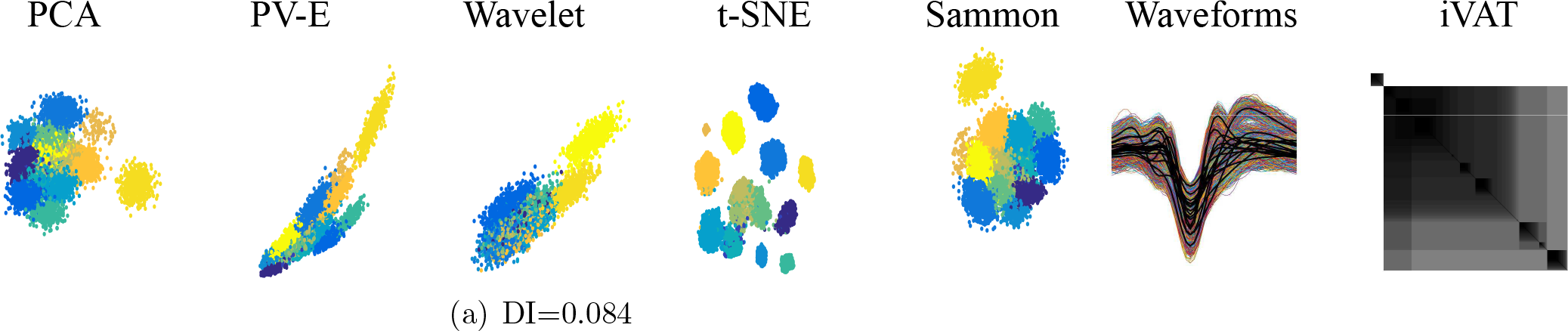
Simulation at c=15. The iVAT image displays 9 blocks and t-SNE shows 13 (colored), and 9 or 10 in black

**Figure 5:**
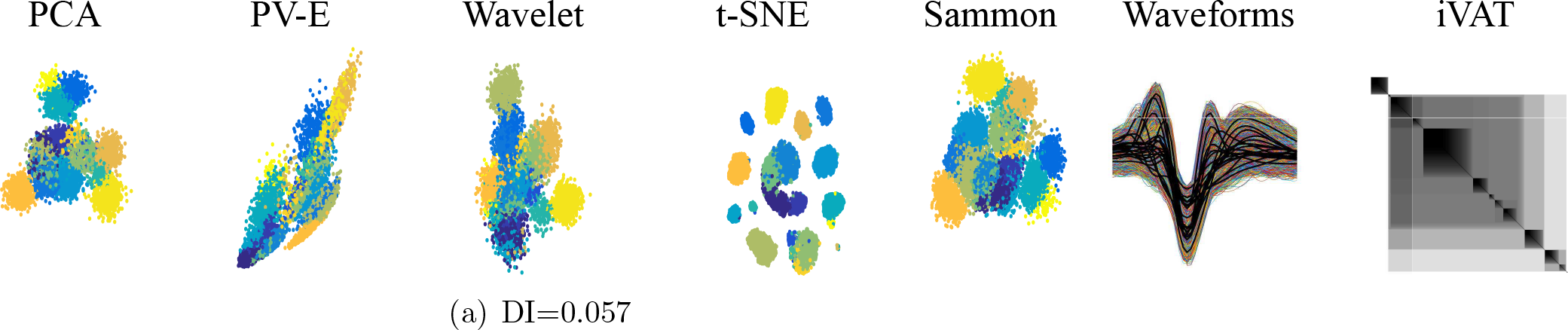
Simulation at c=20. The iVAT image displays 13 blocks and t-SNE shows 12

The next example in this section highlights the ability of iVAT to address two additional problems encountered in spike sorting, namely, anomaly detection and the need for multistage clustering (aka “re-iteration” amongst subclusters). Figure 6(a) is a set called Z of n=4665 waveform vectors comprising a mixture of c=10 labeled subsets from simulated dataset-1. The 10 waveforms shown in Figure 6(b) are the average waveforms, 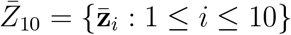, of the ten labeled subsets.

Visual inspection of Figure 6(b) suggests that 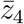, the average waveform of the 488 spikes for unit 4, here called Z_4_, is an outlier (an anomaly) to the general shape of the other 9 graphs. This (easily seen) visual evidence suggests that Z_4_ may form an anomalous cluster in the input or projection spaces. But this observation does not justify removal of all 488 unit 4 spikes from the input data. However, the iVAT image of Z_4_ will corroborate our suspicion that Z_4_ is an anomalous cluster in Z.

Figure 6(c) is the iVAT image of 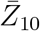 and Figure 6(d) is the dendrogram of the clusters produced by extracting the single linkage hierarchy of clusters from the vectors in 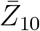. The integers along the borders of the iVAT image of 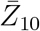 show the identity of each pixel after iVAT reordering. The visualization in 6(c) is quite informative: it not only isolates 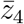 as an outlier (the single pixel at the lower right corner of the image), but it also depicts the other 9 subsets as members of a second large cluster. Moreover, this image suggests a hierarchical substructure within the 9×9 block. The intensities of {5,7} and {6,10} suggest that these pairs of subsets are closely related. The {3,9} block is next in intensity, followed by the 5×5 ping of {8, 5, 7, 9, 3}, which are then coupled to {6,10}, and then this whole structure is embedded within the 9×9 block which includes {1,2}. We remark that the SL hierarchy is easily extracted by applying a back pass that cuts edges in the iVAT MST that reordered 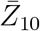 (cf. Kumar et al. [2016]). Figures 6(c) and 6(d) make the relationship between iVAT and single linkage quite transparent. And Figure 6(c) illustrates how an iVAT image can suggest multicluster substructure in a data set.

Figures 6(e) and 6(f) are scatterplots of t-SNE projections of 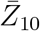 corresponding to perplexity settings of 2 and 3. Both views show the labels of the 10 mean profiles, and both views seem to indicate that 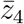 is an outlier in the set 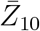. We show these two projections to emphasize that every run of t-SNE with different settings of its hyperparameters may produce different visualizations of its input data. On the other hand, the iVAT image is uniquely determined up to a choice of the distance measure used to construct D.

Figure 7(a) is the iVAT image of the data set Z shown in Figure 6(a). Comparing Figures 6(c) to 6(f) shows that iVAT very clearly suggests the same coarse cluster structure (c=2) in all of the upspace data that it sees in 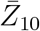, the set of mean profiles. Neither image suggests that c=10; instead, both suggest that the best interpretation of the input data or its means is to first isolate the unit 4 waveform(s), and then regard the remaining spikes as a new cluster, which becomes a candidate for multistage clustering (reclustering, or re-iteration per Niediek et al. [2016]). Note that iVAT makes this information available whether the data are labeled or not.

Finally, Figures 7(b) and 7(c) are labeled and unlabeled t-SNE scatter plots of Z. Both views suggest that Z contains 5 clusters. Subset Z_4_ is isolated in view 7(b), but not more isolated than subset Z_2_, so t-SNE is less assertive about the anomalous nature of Z_4_ than iVAT is. If the labels are available, reclustering might be applied to 9, 7, 5, 8 and/or 3,6,10 to make a finer distinction between spike subsets. If the labels are unavailable, it’s hard to see what can be inferred from the t-SNE projection about Z beyond the suggestion provided by view 7(c) that we should seek 5 clusters in Z.

We conclude this example with some general observations. First, the iVAT image is unique, while t-SNE plots are a function of three user-defined parameters. Second, single linkage clusters of the input data are available via clusiVAT [Kumar et al., 2016] once an iVAT image is built. Third, while Z has 10 labeled subsets of input spikes, neither iVAT nor t-SNE makes this prediction. This emphasizes the fact that labeled subsets may not necessarily be clusters with respect to any computational scheme designed to detect clusters.

**Figure 6:**
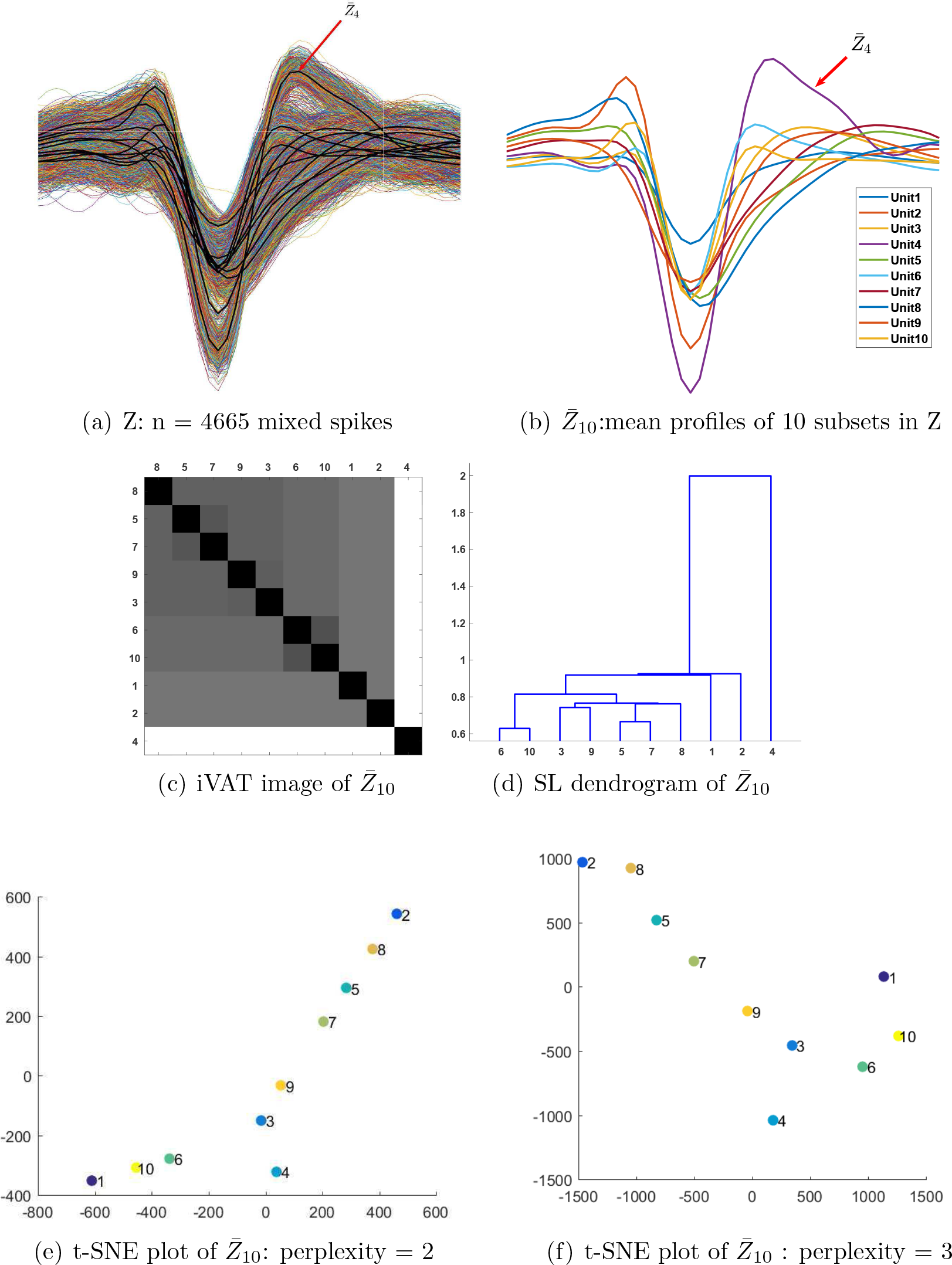
iVAT and t-SNE visualizations of average waveforms of a mixture of 10 subsets of labeled simulated spikes

**Figure 7:**
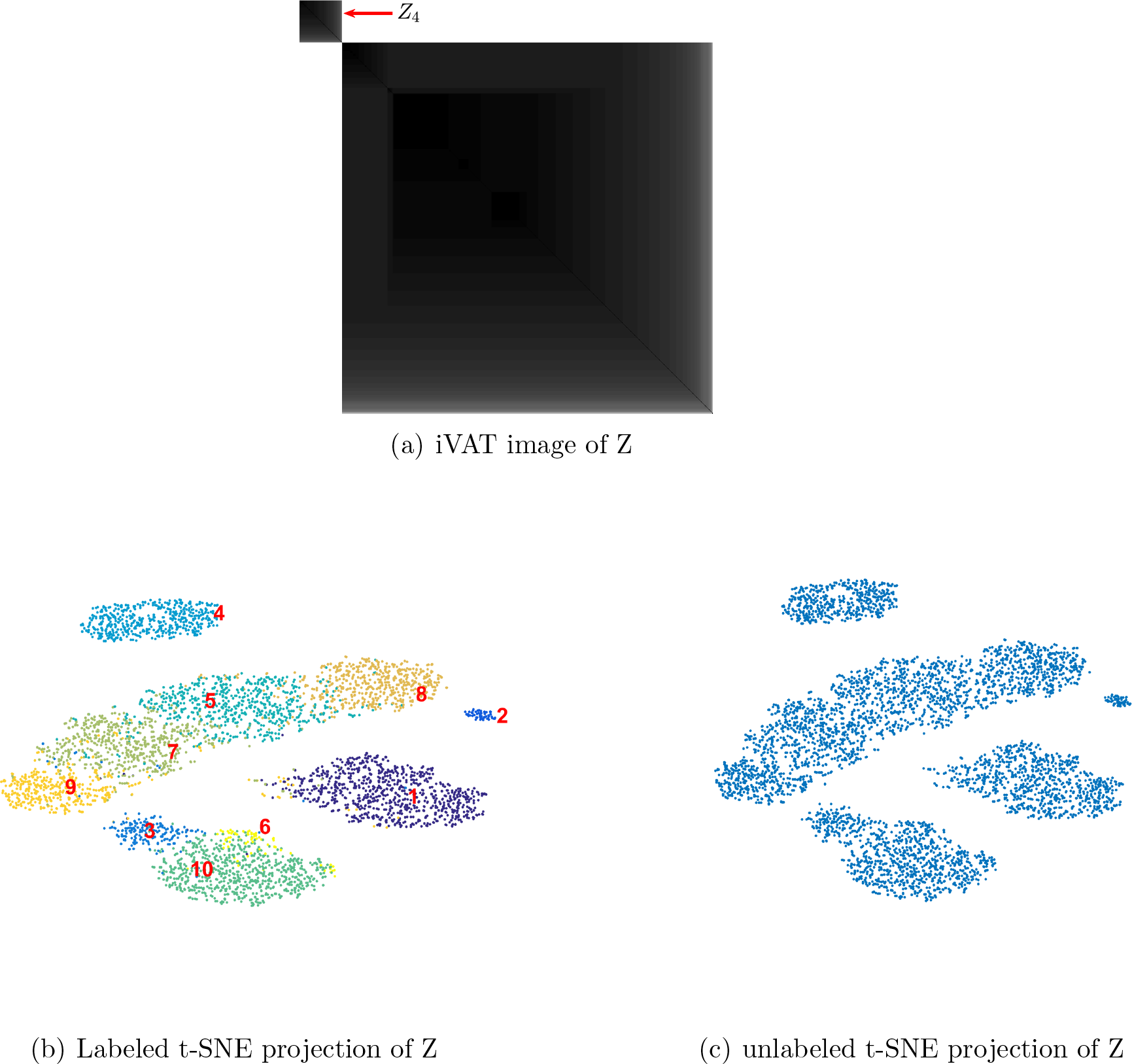
iVAT and t-SNE visualizations of a mixture of 10 subsets of labeled simulated spikes

Now we turn to dataset-2 to further investigate the limits of discernible spike subsets since as mentioned previously, sets drived from dataset-2 are combinations of real spikes originated from pyramidal cells in rat hippocampus (ref to [Henze et al., 2000]). We extracted the spike subsets of nine individual neurons obtained from the different experimental trials. From these nine subsets, we built 36 mixtures at c=2; 84 mixtures at c=3; 126 mixtures at c=4; 126 mixtures at c=5; 84 mixtures at c=6; 36 mixtures at c=7; 9 mixtures at c=8; and one mixture of all nine subsets (c=9). This yields a total of 502 mixtures of labeled waveforms.

For the sake of brevity, we showcase four representative units and the various mixtures that can be built from them at c=2, c=3, and c=4. Figure 8 shows the four representative subsets (all nine subsets of the waveforms are shown in Figure 14).

**Figure 8:**
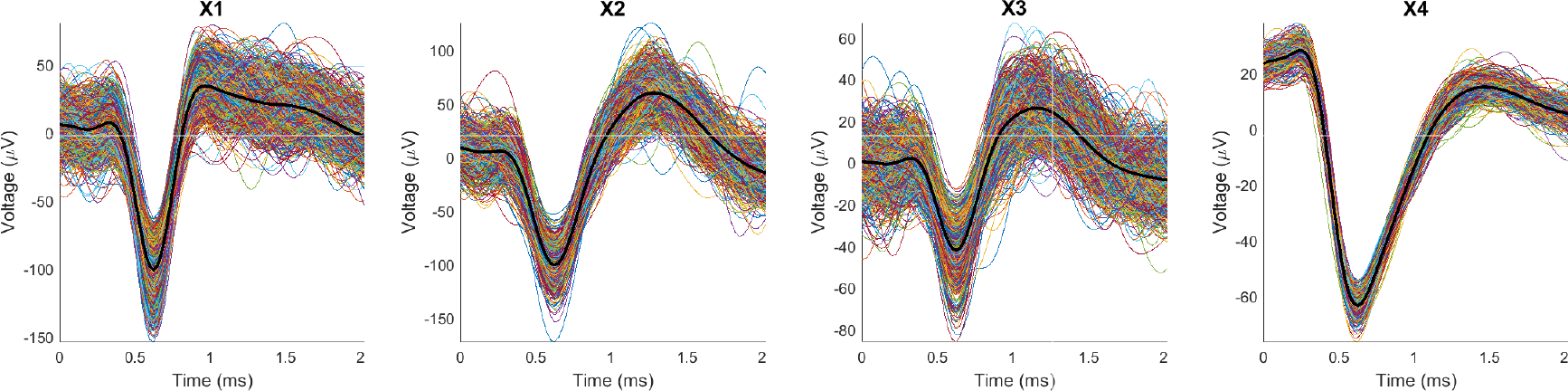
The subsets of spikes generated by four representative units: X1, X2, X3, and X4 containing 1173, 700, 779, and 382 spikes, respectively. Note that the waveforms in X4 are visually different than most of the waveforms in the other three subsets. This fuels an expectation that mixtures with X4 as one component will be somewhat more separable than mixtures without it.

Figure 9 shows all six views of pairs (Xi, Xj) made with 2D transformations of the 80D (upspace, p=80) datasets for the mixtures of two representative single units. We will name the mixtures (Xk, Xj)=Xkj and will follow this convention for all mixtures. For example, the mixture of X1 and X2 is X12, and the mixture of X1, X2, and X4 becomes X124. The waveforms comprising each mixture are also shown, with the average waveform for each single unit in thick black. The colors of points in the 2D scatterplots correspond to class labels of the mixed data. It is important to remember that in a real application, the data are not labeled, so the associated 2D scatterplots will be mono-color dots in the plane. The mixtures are ordered according to increasing values Dunn’s index. Observe that for each mixture, different 2D projections may offer different interpretations of the cluster structure in the upspace data. In 9(a), all five projections show one big cluster, far more evident if the color labeling is missing, which is the case for real experiments in which we do not know the membership of the waveforms. In cases like this, since the clusters are projected densely side by side, human operators or algorithms tend to select only the core of the clusters. This usually produces better values for cluster validity indices, but at the expense of unwarranted confidence in subsequent analyses.

**Figure 9:**
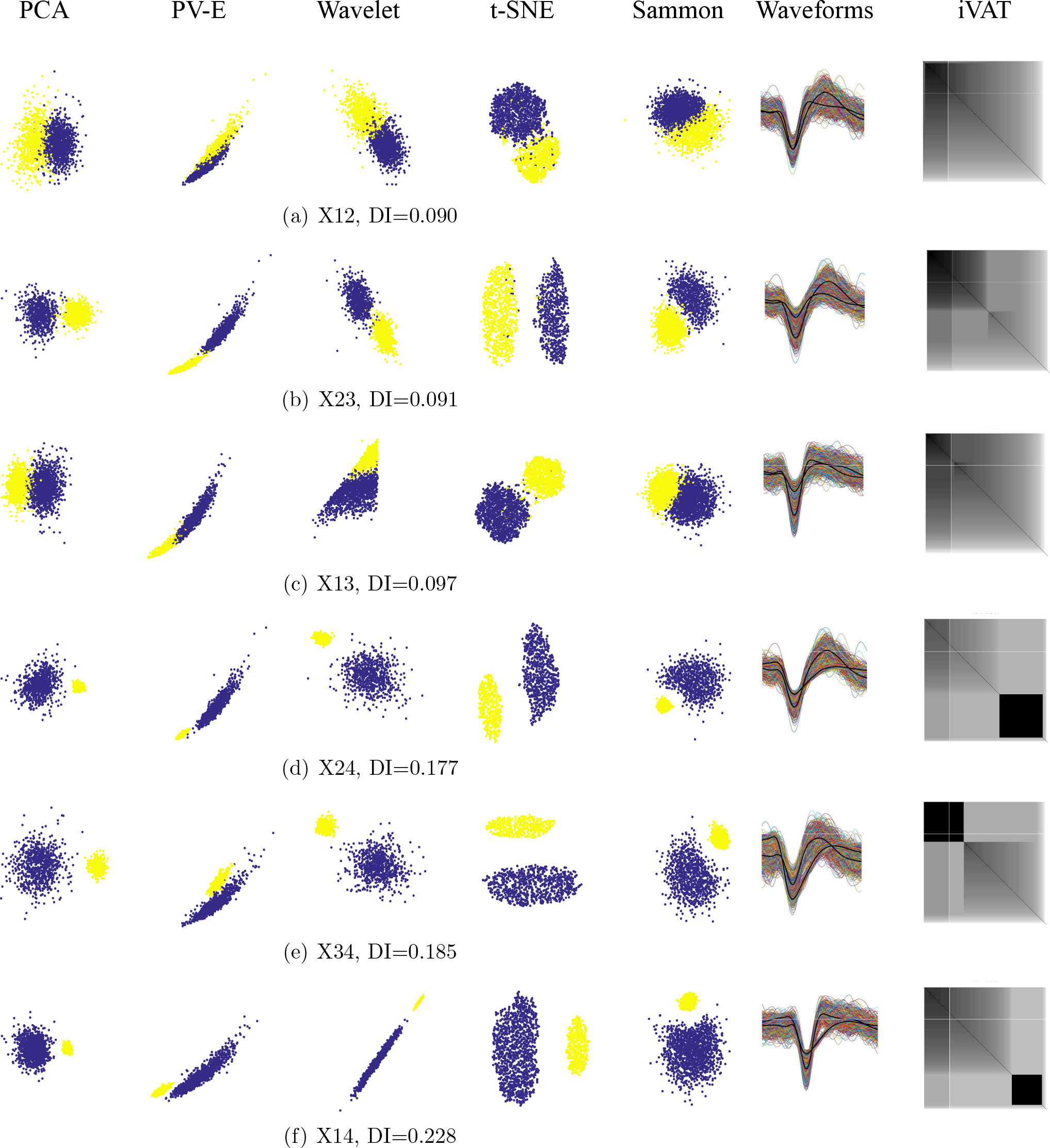
Mixture pairs of X1, X2, X3 and X4, ordered by increasing Dunn’s index

First, some general observations. Figures 9(a), 9(b), and 9(c) all have a DI value of around 0.09. This is a relatively low value that indicates a lack of separation between the two components of the mixture. The iVAT images for these three cases are basically uniform (no strongly visible dark blocks), which indicates that the upspace data are not well separated. Separation emerges in Figures 9(d), 9(e), and 9(f), the three cases that have X4 as one component. Dunn’s index is essentially doubled (0.18 up to 0.23), so upspace separation of the pair of clusters has increased. The most visible separation is seen in the t-SNE downspace scatterplot, which is mirrored in the upspace iVAT images: the strong dark block corresponds to subset X4. Now we will discuss the six cases in more detail.

In 9(b), PCA, Wavelet, t-SNE, and Sammon show two clusters, while the PV-E plot shows just one. In 9(c), PV-E, wavelet, and Sammon projections show one cluster, it can be argued that PCA shows two while t-SNE shows two well-seperted clusters. In the other images, which include X4, a less distorted and noisy set of spikes, all the projections do a good job of mapping the clusters in a separable manner (for the wavelet projection in 9(e), it is hard to see two clusters when there are no color labels). The iVAT image also follows the same trend: the clarity of the two blocks generally becomes higher with a higher Dunn’s index.

The Peak to Valley and Energy (PV-E), are the only real (physically meaningful) 2D features. All the other 2D projections are dimensionless, i.e., they do not have physical meaning. It is important to emphasize that neither the 2D projections nor iVAT produce clusters, all these visual methods just suggest how many to look for.

The projections and the iVAT image of X23 with DI = 0.091 (Figure 9(b)) is a bit more separable and clear than the mixture of X13 with DI = 0.097 (Figure 9(c)). Both values are relatively small, and the difference between these two values (0.008) is negligible, indicating that these two cases are somewhat indistinguishable. The iVAT image for X14 clearly suggests the c=2 at a Dunn Index of 0.228. This provides a much stronger indication of reliability than the smaller DI values. Indeed, Dunn characterized a partition as being compact and separated if and only if DI > 1. DI values less than about 0.5 usually characterize relatively poor cluster structure.

All the cases of mixtures of three subsets are portrayed in Figure 10, again ordered by their Dunn’s index, which is quite low and nearly equal in all four views. The numerator of DI is the minimum distance between any pair of subsets, and the denominator is the largest distance between points in some clusters, so it is dominated by the smallest between-subset distance and largest in-subset distance. Consequently, DI fails to recognize competing clusters that cannot dominate either of the two factors in Dunn’s formulation. These non-dominant clusters can often be seen in iVAT imagery. For example, in Figure 10(b), the small yellow cluster seen in the t-SNE scatterplot of X124 appears as the small dark block in the lower right corner of the corresponding iVAT image. In Figure 10(a) all the projections except for t-SNE fail to point to c=3 and the iVAT image is not informative either. In Figure 10(c) for X234, the PV-E and Wavelet projections suggest that c=2, while PCA, t-SNE and Sammon point to c=3. The t-SNE features provide the widest and most visible separation between the three clusters. The iVAT image of X234 is weakly suggestive of c=3. Figure 10(d) for X134 provides a striking contrast in the ability of the visualization methods to correctly portray the presumed structure in the data. PV-E and Wavelet suggest c=1, PCA and Sammon imply c=2, and t-SNE points clearly to c=3. The iVAT image is pointing to c=2, at a relatively low value of DI=0.097.

**Figure 10:**
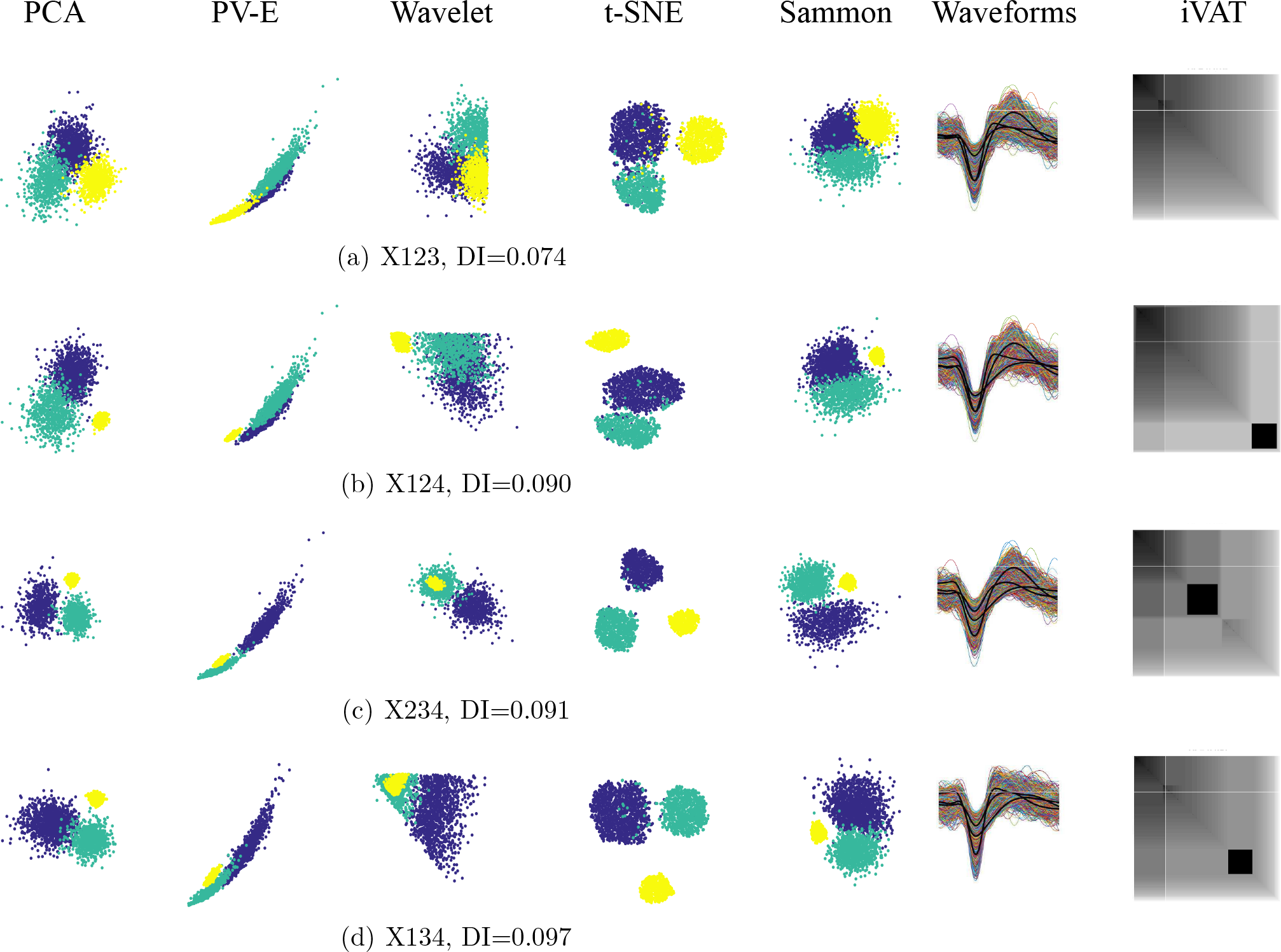
Three-subset mixtures of X1, X2, X3 and X4 at c=3 ordered by increasing values of Dunn’s index

Finally, a similar trend continues in the c=4 subset mixture of X1, X2, X3, and X4. The PV-E and Wavelet features indicate only one big cluster, and PCA, Sammon, and the iVAT image single out X4 while packing the other three sets of waveforms into a single cluster, whereas, t-SNE maps the four subsets with arguably enough clarity to declare that X1234 probably has four clusters. It can be argued that while the input has c=4 labelled subsets, the primary visual evidence does not support c=4, nor will there be a “best” set of clusters in the upspace at this value of c. In other words, just because the subsets have 4 labels does not guarantee that a cluster analysis of the data will agree. When you imagine the scatterplots in Figure 11 without colors there are not four distinguishable clusters present.

**Figure 11:**
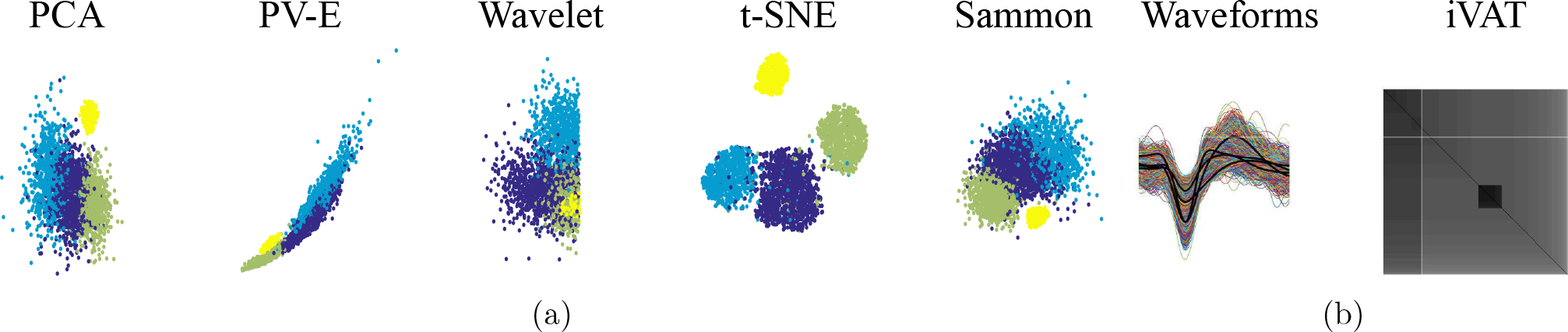
X1234 mixture at c=4

In summary, we note again that although the subsets of spikes were obtained from independent trials, they were all induced by intracellular current injections to hippocampal pyramidal cells and recorded from their close proximity in extracellular medium. This to some degree explains the similar average waveforms and high variability between spikes of one subset. We observed that, for example, from the thirty-six mixtures of two subsets that can be created from the nine spike sets, not all of them were mapped as two clusters with the same quality (i.e. the quality of visualization was not consistent). overall, visualizations of mixtures using any of the other projection methods and iVAT did not suggest discernible clusters. This points out the challenge in identifying neurons of the same class (e.g., pyramidal) from their spike waveforms, at least when they are induced by current injections.

### 3.2 Objective assessment of clustering quality using different projections of the data

So far, we have shown that the lower-dimensional representations in our study may give different interpretations of the upspace data. This problem is highly dependent on the definition of similarity between spike waveforms of different units. Overall, iVAT and t-SNE were most helpful in assessing the pre-cluster presumptive structure of the waveform mixtures. In order to provide a more quantitative assessment of the effectiveness of the different low-dimensional representations in processing spike waveforms, we ran the k-means clustering algorithm on each of the 95 mixtures from dataset-1 and the 502 mixtures from dataset-2.

Dunn’s index and its generalizations provide measures of the intrinsic quality of the computed clusters (based on their distribution with respect to each other). Figure 12 shows the average Dunn’s index (DI) and generalized Dunn’s index (GDI_33_) of the mixtures for the two datasets.

**Figure 12:**
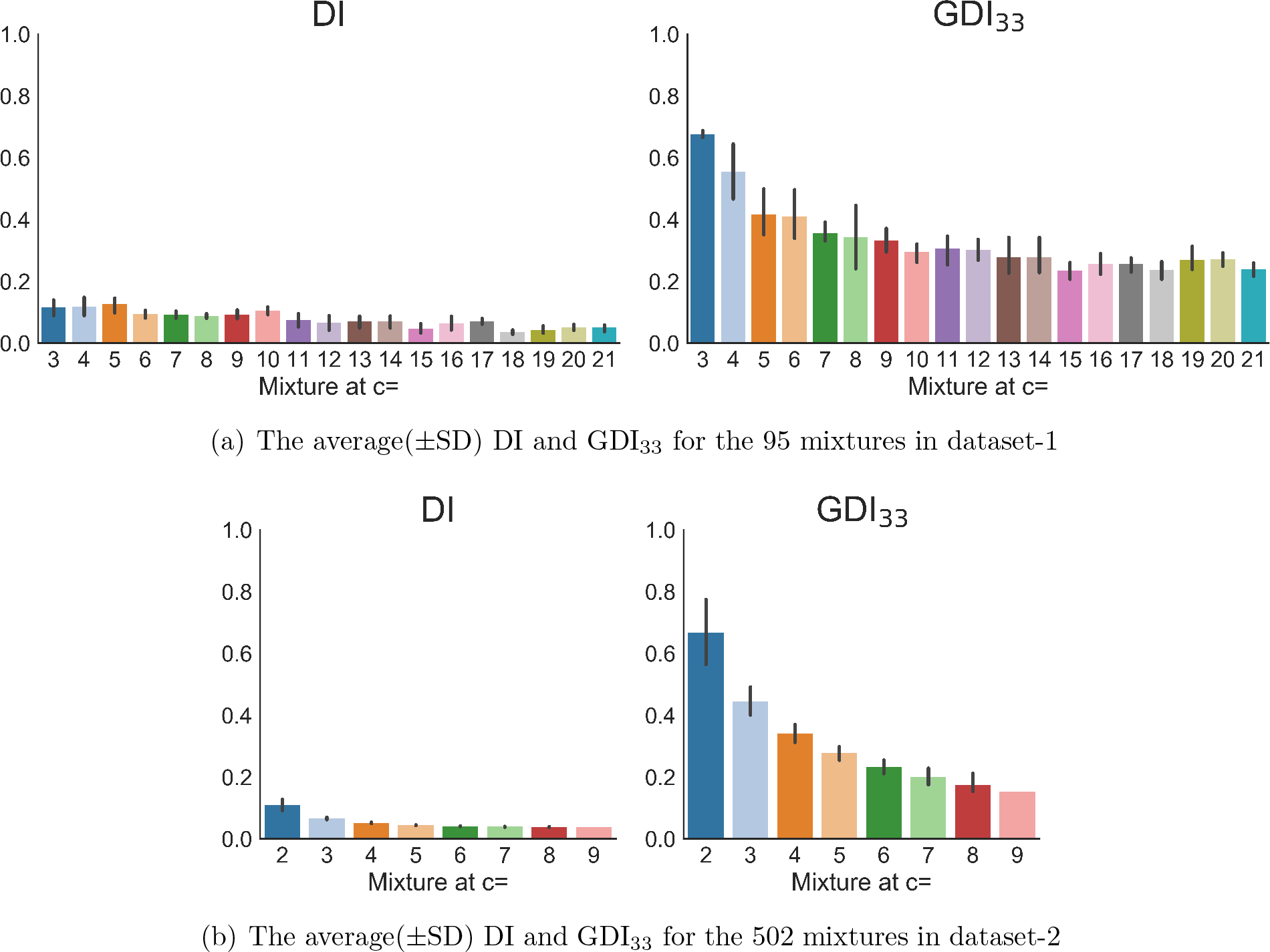
The average(±SD) Dunn’s and generalized Dunn’s indices of ground truth partitions for the mixtures in the two datasets

The two indices have the same trend: they decrease almost monotonically as the number of components (c) increases. However, the generalized version, GDI_33_, provides a much clearer idea of the trend than DI because it has higher values that reflect separation more clearly, and it avoids the bias of inliers and outliers that may affect Dunn’s index. On the other hand, Figure 12 also suggests that both indices tend to favor lower numbers of clusters. This is a different type of empirically observed bias that must be accounted for when relying on cluster validity indices. See Lei et al. [2017] for a discussion related to this point.

The DI and GDI_33_, as internal measures, were used to give a sense of the structure inherent in ground truth partitions of the data in the upspace. Then, to evaluate candidate partitions produced by k-means in the upspace and downspace data sets, we used the adjusted rand index (ARI), which compares the cluster structure of each k-means partition to its ground truth partner at every value of c.

The k-means clustering algorithm is executed on each pair of features obtained with the five methods. The number of clusters to be generated (i.e., c) is set equal to the number of labeled subsets in the ground truth partition (i.e., 2, 3, 4,·. to 9 for cases in dataset-2 and 3, 4, 5, … to 21 for cases in dataset-1). For each computed partition, we calculate the ARI measure of agreement between the computed waveform memberships and the memberships as given by the ground truth partition (recall that our data is labeled).

Figures 13(a) and 13(b) report the average ARI for mixtures in dataset-2 and dataset-1, respectively. In each figure, the first column is the ARI of the clusters achieved by running k-means on the input dimension space (the 48D waveforms for Dataset-1 and the 80D spike waveforms for Dataset-2). The next columns show the average ARIs calculated for the clusters achieved by k-means clustering on the 2D datasets produced by the five techniques. The ARI maximizes at 1, so clustering in the 2D t-SNE downspace data provides k-means clusters that, on average, slightly better match the ground truth partitions than k-means clusters in the input space.

In order to highlight the importance of dimensionality reduction and feature extraction techniques (the pre-clustering stage), this subsection presented a comparison between clustering in the different downspaces and also the input space, using the same clustering algorithm in all spaces. It is important, however, to recognize that the choice of clustering algorithm also contributes to the accuracy of membership assignments. Also, the techniques and parameter choices applied in pre-processing of the signals such as in filtering and spike detection phases alter the end results. Here, we used the classic k-means clustering algorithm as opposed to common spike sorting softwares used by neuroscience community to bypass the different pre-processing techniques that may have affected the down-space representation of spikes. In this way, we provided a controlled set of spikes to be visualized by the 6 methods in our study. Given the good estimate for c provided by visualization with t-SNE projections, a logical next step would be be to substitute this t-SNE features with the one used within one of the softwares and compare the results. We further note that the choice of 2D as opposed to 3D in our manuscript was for publication purposes. Using 3D scatterplots may provide more information that improves the visualization and clustering, but this does not address the “best features” or “best algorithm” issues that follow pre-clustering assessment. Those issues will be addressed in a followup paper.

**Figure 13:**
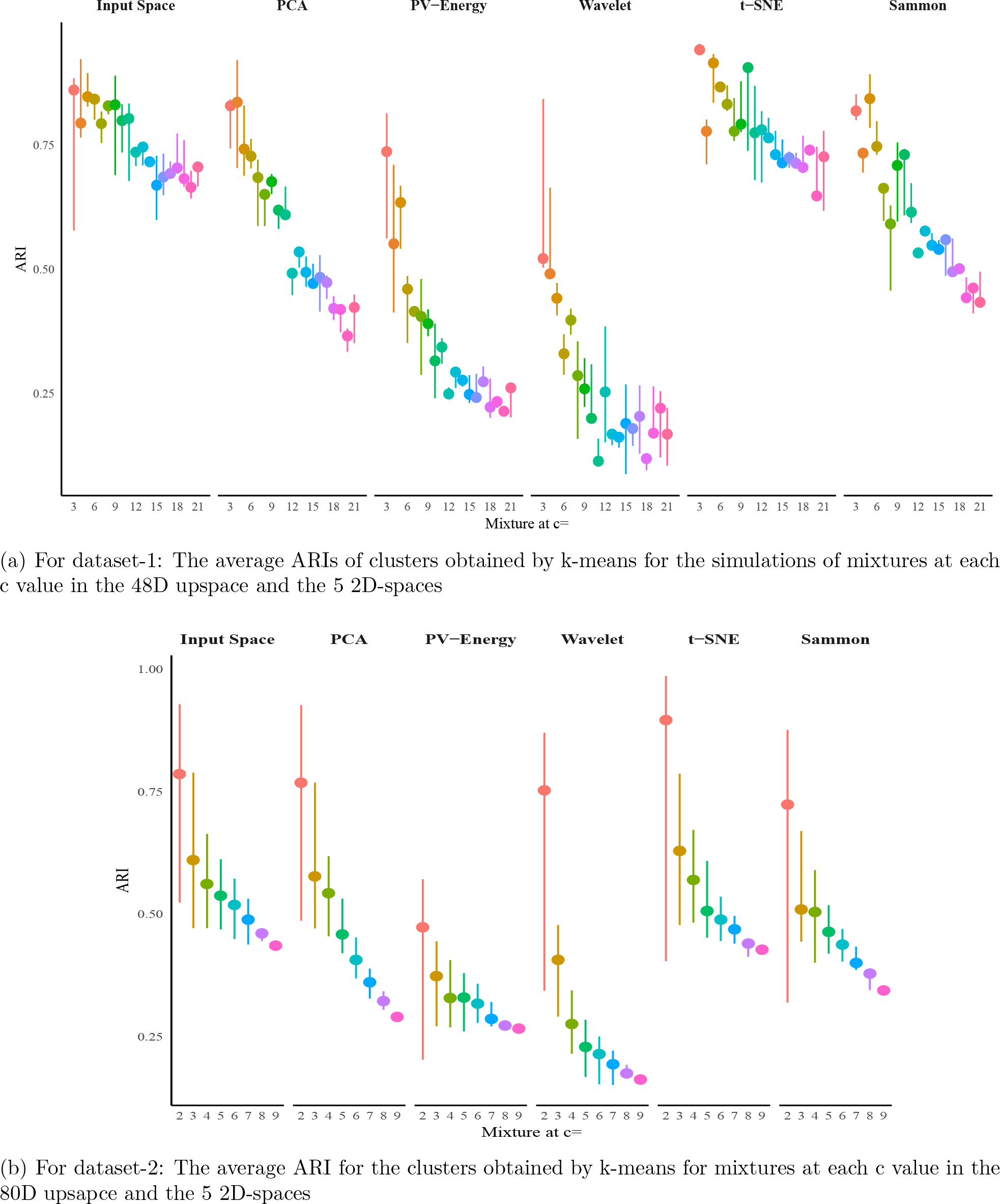
Average validity index (Adjusted Rand index) of the clusters obtained by kmeans on the two datasets

## 4 Discussion and conclusions

Spike sorting is an unsupervised learning problem since the number of neurons associated with the recorded spikes are unknown. Hence, what we consider to be the spike train of a single unit may, in fact, be the spike train of multiple units. This does not negate the usefulness of the previous findings which apply the results of these sorting algorithms to infer neural coding and brain function. Most of this research is based on the rate coding principle [Dayan and Abbott, 2001], which uses the spike rate of the sorted units to model the neuronal response. Rate coding models neglect sorting errors assuming that as long as the spike rate changes according to the stimulus, the model will capture the response, whether the spike train consists of spikes of one or multiple neurons. However, the real units’ tuning curves might be different and a contaminated tuning curve may give misleading results.

This has been shown instudies concerned with issues arising from sorting quality on the results of rate coding models. For example, Todorova et al. [2014] evaluated the quality of the off-line reconstruction of arm trajectories from electrode array recordings and showed that discarding spikes substantially degrades the decoding of the movement to the extent that decoding the unsorted recordings reached higher performance results. They also showed that adding the tuning model (temporal features) of the spiking to the sorting process does not always improve the sorting based on waveform features. We can use the analogy of a verbal fight or discussion among a few people. An observer can tell if the discussion is going smoothly or if it is heated based on the overall volume of the voices of the group, even if the words uttered by individuals is not discernible. This is why rate coding models are popular and successful in certain respects, but they cannot elucidate how neurons interact to give rise to brain functions [Akam and Kullmann, 2014, Huxter et al., 2003, Mehta et al., 2002, Rullen and Thorpe, 2001, Zuo et al., 2015]. This emphasizes that reliable spike sorting is much more critical if any temporal or multiplex coding schemes are to be used to infer neural responses.

Current spike sorting packages use variety of clustering techniques all of which can benefit from an insight to the cluster structure in the data: k-means, Gaussian mixture decomposition or similar algorithms need explicit specification of the number of neurons to seek; mean shift needs the threshold on the bandwidth parameter to be defined; [Comaniciu and Meer, 2002] and density based clustering relies on a threshold on density parameter *γ* for the number of density centers[Rodriguez and Laio, 2014]. In high-dimensional data, the role of visualization in gaining knowledge of the data structure is critical. There is no doubt, as in Plato’s allegory of the cave, that there is always a loss or distortion of structural information in any transformation from the upsace (aka: input space or input dimension) to any downspace. We investigated this issue using iVAT, a tool that enables direct visualization of cluster structure in the upspace as well as five dimensionality reduction methods, including t-SNE. We showed that better sorting can be achieved by securing a visual assessment prior to clustering which affords an estimate of the cluster structure of the data (i.e., the number of clusters, c), or at least a small interval of integers that presumably bracket the true (meaning most distinguishable by some clustering algorithm) but unknown number of clusters.

Our examples show that t-SNE is one of the best methods for projection of high dimensional data to the viewing plane. We note that t-SNE for the present analysis was parameterized with a perplexity of 30 and learning rate of 500. This was the empirically optimized setting for our data and we acknowledge that the need for parametrizing based on the data is a downside to using t-SNE to provide projected data for clustering. The Barnes-Hut variant of t-SNE (publicly available through GitHub) highly accelerates the computations[van der Maaten, 2014]. Since this study focused on investigating the fundamental relevance of pre-clustering methods using standard ground truth datasets the computation time was not substantial to report. Dimitriadis et al. [2016] have a preprint that reports computation speeds of hours for big datasets containing > 10^5^ spikes.

In this paper, we demonstrated that the visual assessment of *c* from the iVAT images is often possible, highlighting that if clustering in the upspace is preferred, a visualization tool such as iVAT can be integrated into the package to inform the manual curation process. However, there were cases, in particular when *c* was high (> 10), that the iVAT image did not clearly indicate the number of clusters. But the visual assessment of a user that makes the estimate of *c* subjective. We showed that extracellular neuronal waveforms generate noisy datasets that at times do not comprise well-separated clusters. So, iVAT does not provide the definitive answer to the problem of spike sorting. Nevertheless, it provides insight into the coarse structure of the dataset. Moreover, we mentioned the relationship between iVAT and the single linkage clustering algorithm that is illustrated in Figures 6(c) and 6(d) (See Havens et al. [2009] and Mahallati et al. [2018a] for further discussion). The majority of the edges in the MST that iVAT builds connect neighbor points and hence have very small values. The largest values in the MST usually correspond to edges that connect clusters (and the outlier points). The threshold between the small and large values reflects finer distinctions between clusters in the upspace, which can be used in assigning spikes to clusters. This feature was integrated into scalable and faster implementations of the iVAT algorithm which can be used for big datasets [Rathore et al., 2018].

Our experiments on visualization of the two labeled datasets provided further insights into spike sorting. In the first dataset, simulations were generated using average waveforms obtained from extracellular recordings in behavioral experiments. For the mixtures of spike subsets extracted from this dataset it was possible to estimate the presumptive cluster number in the data from the dark blocks in the iVAT images, even in some cases of mixtures of twelve subsets. Mixtures of higher subsets were sometimes displayed as compact and isolated clusters in the t-SNE projections. Our experiments confirm that when the data possess compact, well-separated clusters, visualization can be quite useful. Dataset-1 represents mixtures of spike sets that are generated by different cell types, brain regions and brain states, and these can be distinguished based on their spike waveforms. In contrast, dataset-2 represents mixtures of spike sets that are induced from cells of the same class receiving intracellular current injections, hence providing spikes with similar waveforms. Therefore, classifying based on extracellular waveforms alone may not be feasible in the latter case (cells of the same type receiving the same input). It should be noted that in sorting of spikes for each electrode, different distances of the units from the electrode improves sorting, since the amplitude (energy) of the waveforms is different. However, these results indicate that a 660 further subtype classification beyond the two main classes of inhibitory and pyramidal categories (i.e. subtypes of pyramidal cells) may not be feasible by considering only the spike waveforms.

The extracellularly recorded potentials are already distorted signatures of intracellular action potentials, which makes the dimensionality reduction stage even more critical. The problem of crowding in lower dimensional maps such as PCA is well known. Analysis of the simultaneous extracellular and intracellular recordings have shown that the probability distributions of spikes from different neurons in the PCA feature space have some degree of overlap [Harris et al., 2000]. There is an inherent variability in the extracellular waveforms imposed by the background field potential [Buzsáki et al., 2012], variations in the intracellular action potentials due to factors such as bursting [Henze et al., 2000], and slight electrode drift over the course of the experiment [Harris et al., 2016]. Activity of neighboring neurons is also a possible source of distortion in the waveform shape. Such activity may sometimes overlap in time and make multi-unit spike waveforms. The problem of overlapping spikes has been addressed by methods such as independant component analysis, ICA, (if number of electrodes is equal to or more than the number of neurons)[Takahashi et al., 2003] or template matching [Yger et al., 2018]. It is worth noting that in pre-clustering visualization of the data, overlapping spikes (multi-unit spikes) construct a cluster or block. Visuliza-tion is not to substitue post-clustering use of ICA or template matching. These methods can be used after clustering to extract individual sources of multi-unit spikes and reassign these spikes to the previously identified clusters. With regards to the inherent variability in the extracellular waveforms, our results show that the t-SNE projection is the most reliable feature extraction scheme (for visualization) that we tested. We believe that t-SNE works well since it is a probabilistic-based approach that is appropriate for neuronal data. In a nutshell, the variability caused by the noisy spikes can often be circumvented by converting the deterministic dissimilarity measure between two waveforms into a probability of dissimilarities.

Another reason why having a reliable dimensionality reduction stage is important is revealed by our results on Dunn’s index, which showed that, DI, in common with many other internal cluster validity indices, tends to be monotonic in *c*. This emphasizes the point that the common practice of running a clustering algorithm for several values of *c* and then choosing the best partition based on the optimal value of any cluster validity index may not be very effective. Moreover, by computing both DI and GDI_33_ for the same data, we demonstrated that there is no agreement about a generic CVI, a fact that has been shown before in previous experiments on internal CVIs [Vendramin et al., 2010]. Indeed, in the real (unlabeled data) case, it is wise to compute a number of different internal CVIs, with a view towards ensemble aggregation of the results. To appreciate the disparity that different CVIs can cause, see [Arbelaitz et al., 2013] for an extensive survey of 30 internal CVIs tested on 20 real data sets. See [Vega-Pons and Ruiz-Shulcloper, 2011] for a survey of ensemble approaches to clustering. Fournier et al. [2016] have applied this method to aggregation of partitions obtained by different clustering methods used for sorting spike waveforms. Here we suggest using an ensemble approach on the votes cast by different internal cluster validity indices - DI and its 18 GDIs are just a few of the ones available in Arbelaitz et al. [2013] - for each partition in CP. We think this approach will greatly improve the final interpretation of structure in unlabeled data. This will be the objective of our next foray into spike sorting clustering algorithms.

This study again confirms that there is no such thing as the best set of features for clustering or the best clustering algorithm for spike sorting, but that sorting is an iterative process that always comprises making a compromise between the best feature set and clustering algorithm. Armañanzas and Ascoli [2015] list the identification of the number of clusters as the most outstanding question in techniques for neuronal classfication. This challenge can be partially addressed by subjective visual assessment of cluster tendency. While visual evidence is never enough, it has great value, as noted by the eminent statistician Sir Ronald Fisher, who said, nearly 100 years ago: “The preliminary examination of most data is facilitated by the use of diagrams. Diagrams prove nothing, but bring outstanding features readily to the eye; they are therefore no substitute for critical tests as may be applied to the data, but are valuable in suggesting such tests, and in explaining conclusions founded upon them”[Fisher, 1958].

## Acknowledgments

This research was supported by the Natural Sciences and Engineering Research Council of Canada. MRP was funded by Dean Connor and Maris Uffelmann Donation, Canadian Fund For Innovation and Natural Sciences and Engineering Research Council: Discovery Grant (RGPIN-2016-06358). We confirm that the preprint of this article has been submitted to BioRxiv [Mahallati et al., 2018b].

## A Appendix Supporting material

## A.1 Supplementary figure

**Figure 14:**
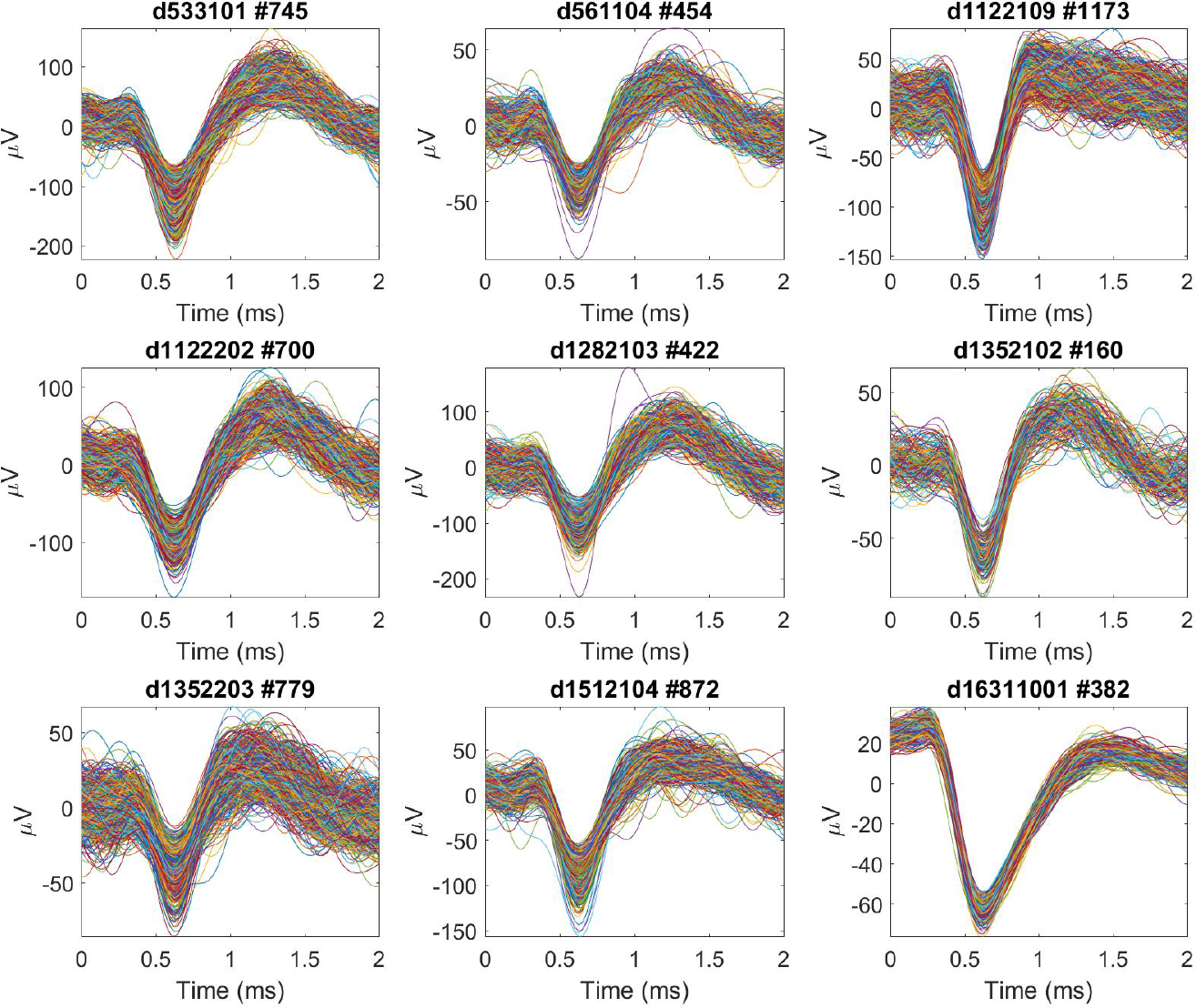
The subsets of spikes of the 9 individual neurons used in the study. Each subplot title displays the label of the experiment in [Henze et al., 2009] dataset and the number of spikes in each subset: e.g., #745 means there are 745 waveforms in the sample.

## A.2 iVAT algorithm

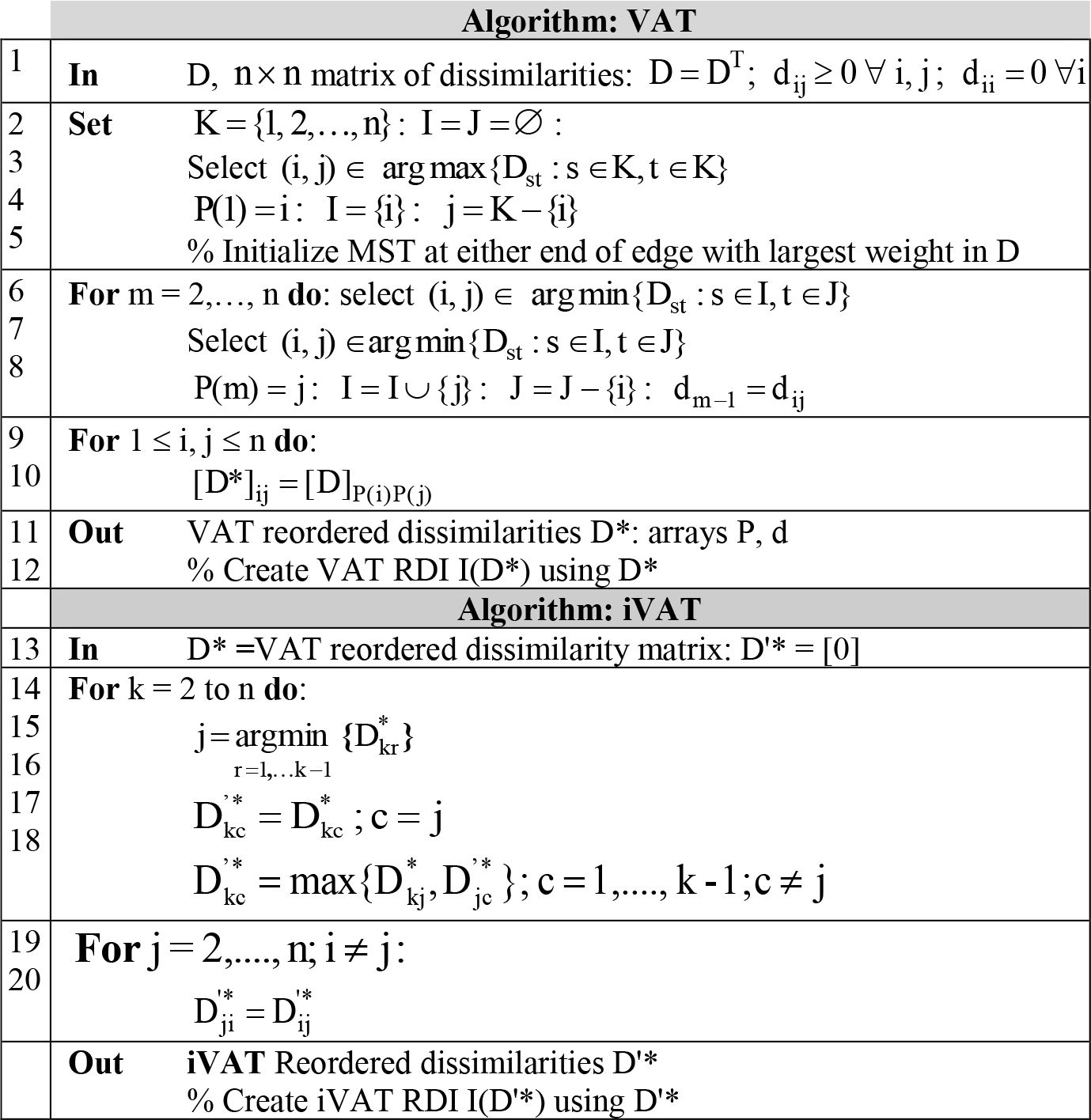

